# Active ribosome profiling with RiboLace

**DOI:** 10.1101/179671

**Authors:** Massimiliano Clamer, Toma Tebaldi, Fabio Lauria, Paola Bernabò, Rodolfo F. Gómez-Biagi, Elena Perenthaler, Daniele Gubert, Laura Pasquardini, Graziano Guella, Ewout J.N. Groen, Thomas H. Gillingwater, Alessandro Quattrone, Gabriella Viero

**Affiliations:** Centre for Integrative Biology, University of Trento, Via Sommarive, 9 Povo (Italy).; IMMAGINA Biotechnology srl, Via alla cascata 56/c, Povo (Italy).; Institute of Biophysics, CNR Unit at Trento, Via Sommarive, 18 Povo (Italy).; Fondazione Bruno Kessler-LaBSSAH, Via Sommarive 18, Povo, Trento, Italy.; Department of Information Engineering and Computer Science, University of Trento, Povo (Italy).; Department of Physics, University of Trento, Povo (Italy).; Edinburgh Medical School: Biomedical Sciences, University of Edinburgh, Edinburgh, UK

## Abstract

Ribosome profiling, or Ribo-Seq, is based around large-scale sequencing of RNA fragments protected from nuclease digestion by ribosomes. Thanks to its unique ability to provide positional information concerning ribosomes flowing along transcripts, this method can be used to shed light on mechanistic aspects of translation. However, current Ribo-Seq approaches lack the ability to distinguish between fragments protected by ribosomes in active translation or by inactive ribosomes. To overcome these significant limitation, we developed RiboLace: a novel method based on an original puromycin-containing molecule capable of isolating active ribosomes by means of an antibody-free and tag-free pull-down approach. RiboLace is fast, works reliably with low amounts of input material, and can be easily and rapidly applied both *in vitro* and *in vivo*, thereby generating a global snapshot of active ribosome footprints at single nucleotide resolution.

---

The tightly regulated process of protein synthesis is a core regulator of numerous critical physiological pathways, ranging from cell growth^1^ and development^2,3^ through to immune responses^4^. Local protein synthesis in neurons^5^ also plays fundamental roles in memory formation^6–8^ and synaptic plasticity^9^. Hence, dysregulation of translation is a major driver of important pathologies, such as cancer^10,11^ and neurodegenerative diseases^12^.

Over the last few years, newly developed methodological approaches such as ribosome profiling (Ribo-Seq)^13^, have contributed to considerable new insights into the translation process. Ribo-Seq has been largely employed to identify novel translated RNAs (coding and non-coding), map novel upstream Open Reading Frames (ORFs), and estimate translation levels in different biological conditions. Indeed, Ribo-Seq has the potential to estimate translation efficiencies and the “protein synthesis levels”^14,15^ in a variety of organisms, from prokaryotes^15^, to yeast^13^, *C. elegans*^16^, zebrafish^17,18^, plants^19^, the mouse^20^, and human cell lines^21–23^. Despite its undoubtful discrimination power and wide applicability, Ribo-Seq still faces a number of challenges and presents with numerous limitations. For example, translationally inactive mRNAs can be sequestered into ribonucleoprotein particles (mRNP) or monosomes (80S), whose translational status remains a controversial issue^24–26^. Importantly, mRNAs can be trapped within stalled or paused polysomes, as has been shown especially in neurons^12,27–33^. As such, Ribo-Seq does not necessarily discriminate “true” protected footprints of translating polysomes from RNA fragments protected by the 80S ribosome or stalled ribosomes in polysomes, leading to possible misinterpretations of translation occupancy profiles. Therefore, Ribo-Seq still requires further optimization and refinements, from the bench to the data analysis^34^, in order to generate maximal insights into the translation process. Here, we present RiboLace, a novel methodological approach based on a newly developed reagent, a puromycin analog molecule. RiboLace greatly improves the study of translation both *in vitro* and *in vivo*, by selectively portraying mRNA fragments protected by *bona fide* active ribosomes at single nucleotide resolution and with unprecedented simplicity, requiring 30 times less biological material than current protocols.

## RESULTS

### Design and synthesis of a new analog of puromycin

Puromycin is an aminonucleoside antibiotic able to bind the catalytic center of the ribosome and the nascent peptide chain, causing ribosome disassembly and disruption of protein synthesis^35–38^. Over the years, it has been used extensively to quantify global of protein synthesis, taking advantage of radioactive^39^ and biotinylated molecules^40^ or anti-puromycin antibodies^41^. Leveraging its ability to keep contact with the ribosome^42–45^, puromycin has also been employed to covalently link an mRNA to the corresponding protein during its synthesis^46,47^. In addition, puromycin can be modified to create cell-permeable analogues suitable for direct and *in situ* imaging of newly synthesized proteins^48,49^. All these methods require the irreversible reaction of the α-amino group of puromycin with the carbon on its carbonyl group, acylating the 3’ hydroxyl group of the peptidyl-tRNA buried in the P-site of the ribosome.

Motivated by evidence that molecules containing α-amino group modified puromycin can bind the large subunit of the active ribosome^45^, we covalently coupled puromycin to a biotin moiety through two 2,2’-ethylenedioxy-bis-ethylamine units, to obtain a new compound still able to bind ribosomes by mimicking the tRNA entrance in the acceptor site (A-site). We synthesized the new molecule at purity higher than 90%, characterized it by NMR (Fig. 1a and Supplementary Fig. 1-6) and called it 3P. Then, we verified the activity of the biotin moiety, taking advantage of its absorbance spectrum and tested the binding on polystyrene or agarose beads. We observed that the biotin group allows the binding of the 3P to commercially available streptavidin-beads (Fig. 1b).

**Figure 1.**
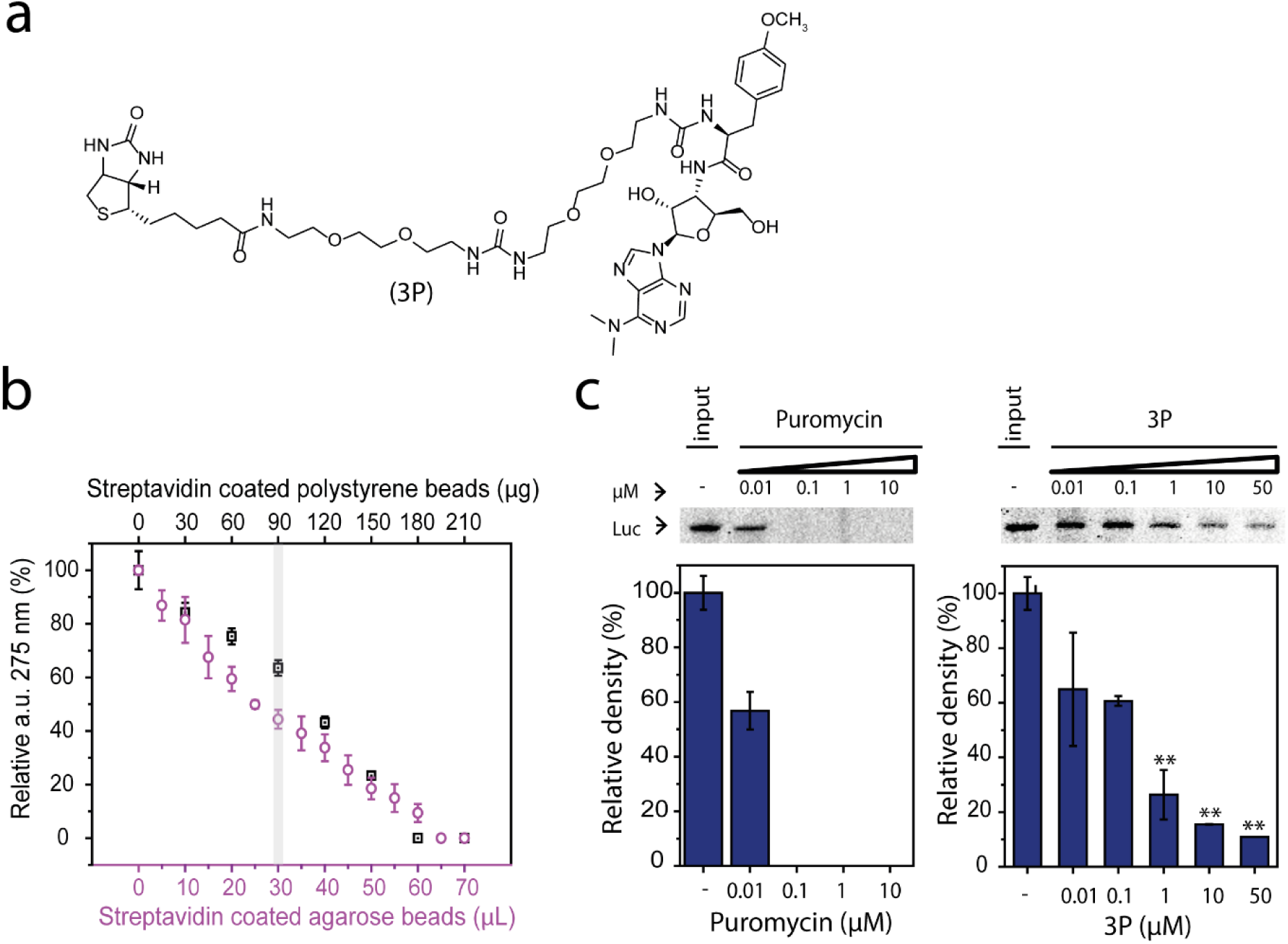
A new analog of puromycin inhibits translation and can be used for functionalization of agarose and polystyrene beads. (**a**) Chemical formula of the new puromycin analog: 3P. (**b**) 3P binding to streptavidin coated agarose and polystyrene beads. Absorbance of the supernatant at 275 nm is measured after addition of streptavidin coated magnetic beads to 100 pmol of 3P. Data represent the mean of triplicate experiments (n = 3). The gray line identified the quantity of beads (polystyrene, μg or agarose, μL of 10% slurry suspension) used in all experiments for each sample. (**c**) Expression of the firefly luciferase in the presence of puromycin (left panel) and 3P (right panel). ε-Labeled biotinylated lysine-tRNAs is used to monitor the protein production by SDS–PAGE (top). The histograms represent the relative quantification of the representative bands reported in the immunoblot with respect to the control. The histogram on the right shows inhibition of luciferase expression in the presence of different concentrations of 3P (0 μM, 0.01 μM, 0.1 μM, 1 μM and 10 μM (right panel). Error bars represent s.d. calculated from triplicate experiments (n =3); (**) = t-test p-val < 0.05.

To demonstrate that 3P molecule maintains an inhibitory effect on translation, we compared its effects to that exerted by puromycin using an eukaryotic *in vitro* cell-free transcription-translation system (IVTT, rabbit reticulocyte lysate) and the firefly luciferase as a reporter gene. We monitored total protein production by SDS-PAGE (**Fig.1c, Supplementary Fig.7**) and luminescence assay **(Supplementary Fig.7**) in the presence of puromycin and 3P at different concentrations. As expected, puromycin induced conspicuous decay of protein production at nanomolar concentrations. In the case of 3P, we observed a decreased level of translation, which reached ∼ 70% of inhibition at concentrations higher than 1 μM (Fig. 1c). We concluded that, even if with slightly lower efficiency than puromycin, 3P can inhibit eukaryotic translation *in vitro*, interfering with ribosome function, and can be used to produce functionalized beads.

### 3P-functionalized magnetic beads captures mRNAs in active translation *in vitro.*

Our finding that 3P is able to interact with the translation process, likely through its puromycin moiety, prompted us to investigate whether 3P can capture mRNAs under active translation.

First, we monitored the ability of 3P-functionalized magnetic beads (3P-beads) to purify transcripts of reporter genes with different levels of protein expression in *in vitro* translation systems. To purify mRNAs with 3P-beads, we developed the following protocol (Fig. 2a): (i) 3P-beads and control beads functionalized with a biotin-glycol conjugate (mP-beads, see Fig. 2a legend and methods for details) were added to the *in vitro* translation system and the suspension was incubated for one hour at 4°C in an appropriate buffer containing cycloheximide, to antagonize the dissociation of ribosomal subunits exerted by the puromycin component of the molecule; (ii) beads were pulled down using a magnet and washed two times; (iii) protein and/or RNA were extracted for downstream analyses. In parallel, the production of protein was followed by immunoblotting of whole protein extracts.

**Figure 2.**
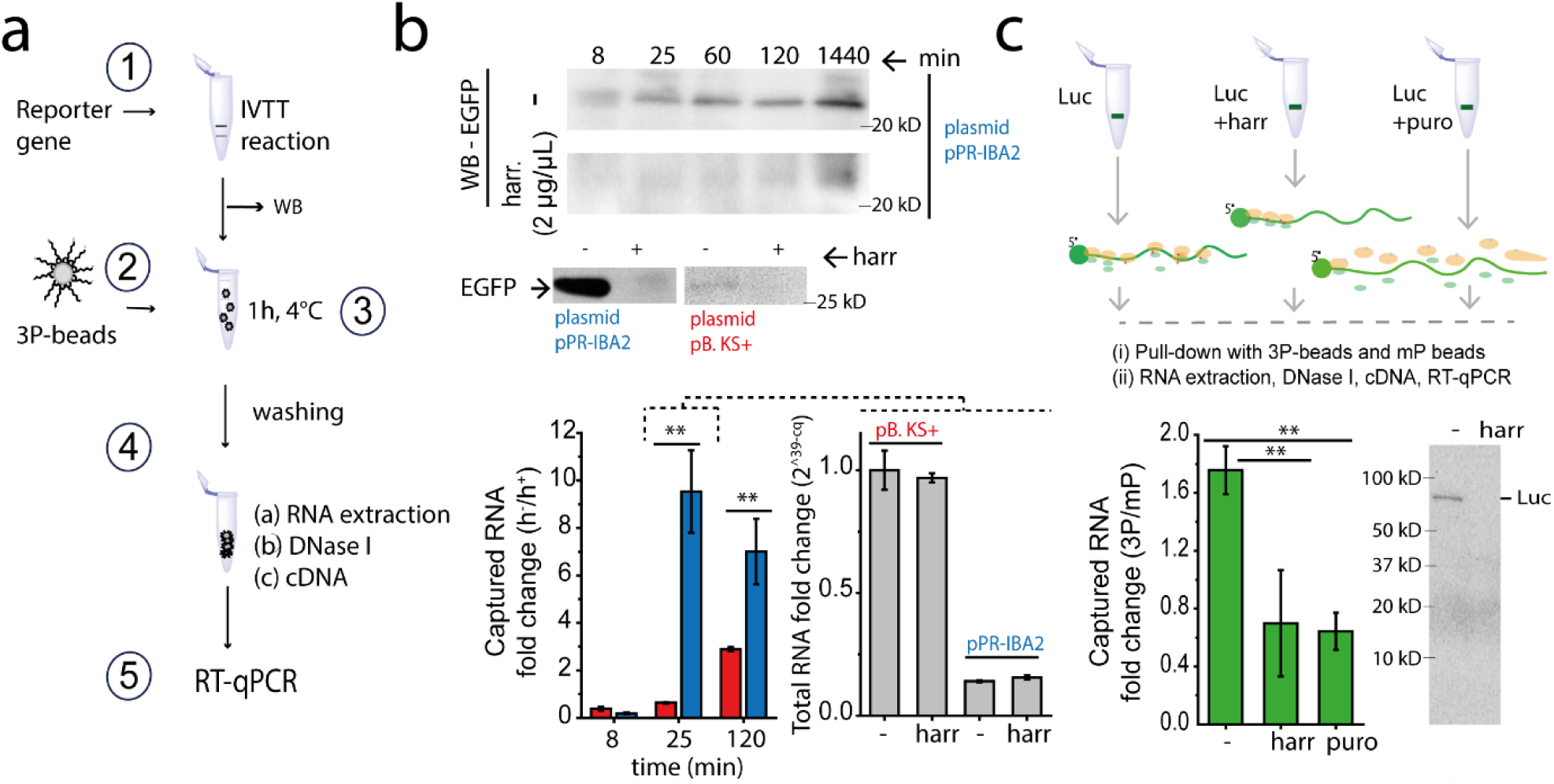
3P-beads can capture mRNAs in active translation *in vitro*. **(a)** Experimental design: 3P-nbeads are used to pull-down transcripts in a cell-free in vitro transcription-translation system. Briefly, from step 1 to 5: (1), Plasmids are added to the *in vitro* transcription-translation reaction; (2), Beads are functionalized with 3P; (3), The IVTT mix is added to 3P-beads and incubated for 1 hour in orbital rotation at 2 rpm at 4°C; (4), Beads are washed without detaching them from the magnet to remove unspecific binding; (4/a), The RNA is extracted, precipitated with isopropanol, and digested with DNase I (4/b) to avoid possible DNA contaminations. Finally, the cDNA is synthetized (4/c). (5), Samples are analyzed by RT-qPCR to detect the reporter gene. **(b) Top panel**, immunoblotting of total EGFP protein at different incubation time, without (-) or with harringtonine (+) at the reported concentration. **Middle panel**, immunoblotting showing the comparison between the total EGFP expression from the pPR-IBA2 plasmid and the EGFP expressed from the pBluescript II KS+ plasmid. **Bottom panel**, EGFP RNA enrichment on RiboLace (-, no harrigtonine) respect to the treated sample (harr, harringtonine treatment, 2 μg/mL for 3 min). On the right, the the total RNA content (gray histograms) without (-) or with harringtonine (harr) for the two vectors as measured by RT-qPCR. (**) = t-test pval = 0.01 (25 min); pval = 0.03 (120 min); n = 3. **(c) Top panel**, Sketch of experimental protocol for in vitro transcription and translation of the firefly luciferase(luc). Harringtonine and puromycin (puro, 5 μg/mL) are added to the mixture. Then, (i) ribosomes in active translation are isolated with 3P-beads, by using mP-beads as control; (ii) RNA is extracted, treated with DNAse I and retro-transcribed to single stranded cDNA with random hexamers. **(c) Bottom**, Fold change values relative to the total amount of transcript captured by 3P-beads compared to the control beads (mP), henceforth referred as the ’enrichment’. (**) = t-test pval = 0.03 (luv vs luc+harr), pval = 0.02 (luc vs luc+puro); n = 5. Error bars represent s.d.

As expected, the EGFP reporter gene showed differential efficiency in protein production depending on the expression vector used (pBluescript II KS+^50^, low performance protein production and IPR-IBA2^51^, high performance protein production) (**Fig. 2b, upper panel and Supplementary Fig. 8**). Complete inhibition of protein production was observed after addition of the translation inhibitor harringtonine, as a control. RNA was purified and the relative abundance of the EGFP mRNA, in both low and high performant conditions, was monitored by qRT-PCR. We observed on 3P functionalized beads a 7 to 10-fold enrichment of the reporter transcript in conditions of active translation, with respect to samples treated with harringtonine (Fig. 2b **lower panel, left**), and in the absence of transcriptional changes (Fig. 2b **lower panel, right**). The decreased enrichment in highly performant conditions and at 120 min of incubation can be related to the fact that the *in vitro* system is a closed one, causing the reaction to stop in the absence of continuous supply of reactants during later stages of incubation.

Then, to demonstrate that this result was not dependent on the reporter used, we applied RiboLace to a luciferase reporter system. In this case we observed a > 1.6-fold enrichment of luciferase transcript with respect to negative controls (mP-beads) and an enrichment with respect to samples where translation had been inhibited (Fig. 2c), as previously observed for EGFP (**Supplementary Fig. 9**). Finally, to understand if the observed enrichments were dependent on the puromycin moiety of 3P, we pre-saturated the system with puromycin and induced ribosome drop-off. Under these conditions, we found no evidence for mRNA enrichment. Overall, these findings support the claim that 3P-beads can be used to capture transcripts undergoing translation in eukaryotic *in-vitro* systems. We named the method RiboLace.

### RiboLace captures active ribosomes and associated mRNAs from whole cellular lysates

Next, we wanted to establish whether RiboLace was capable of isolating ribosomes and mRNAs under active translation from more complex samples than *in vitro* mixtures. We used RiboLace on whole cellular lysates and under different translational states, and monitored the resulting proteins and mRNAs associated with the beads (Fig. 3a). We took advantage of established cellular stimuli that can induce cells into translationally active or inactive states. To shut-down translation we used cell starvation, oxidative, proteotoxic and heat stresses, known to globally suppress protein synthesis^52^ (**Supplementary Fig. S10a**). To specifically activate protein synthesis, we rescued cells from starvation by Epithelial Growth Factor (EGF) or by Fetal Bovine Serum (FBS) stimulation^53^.

**Figure 3.**
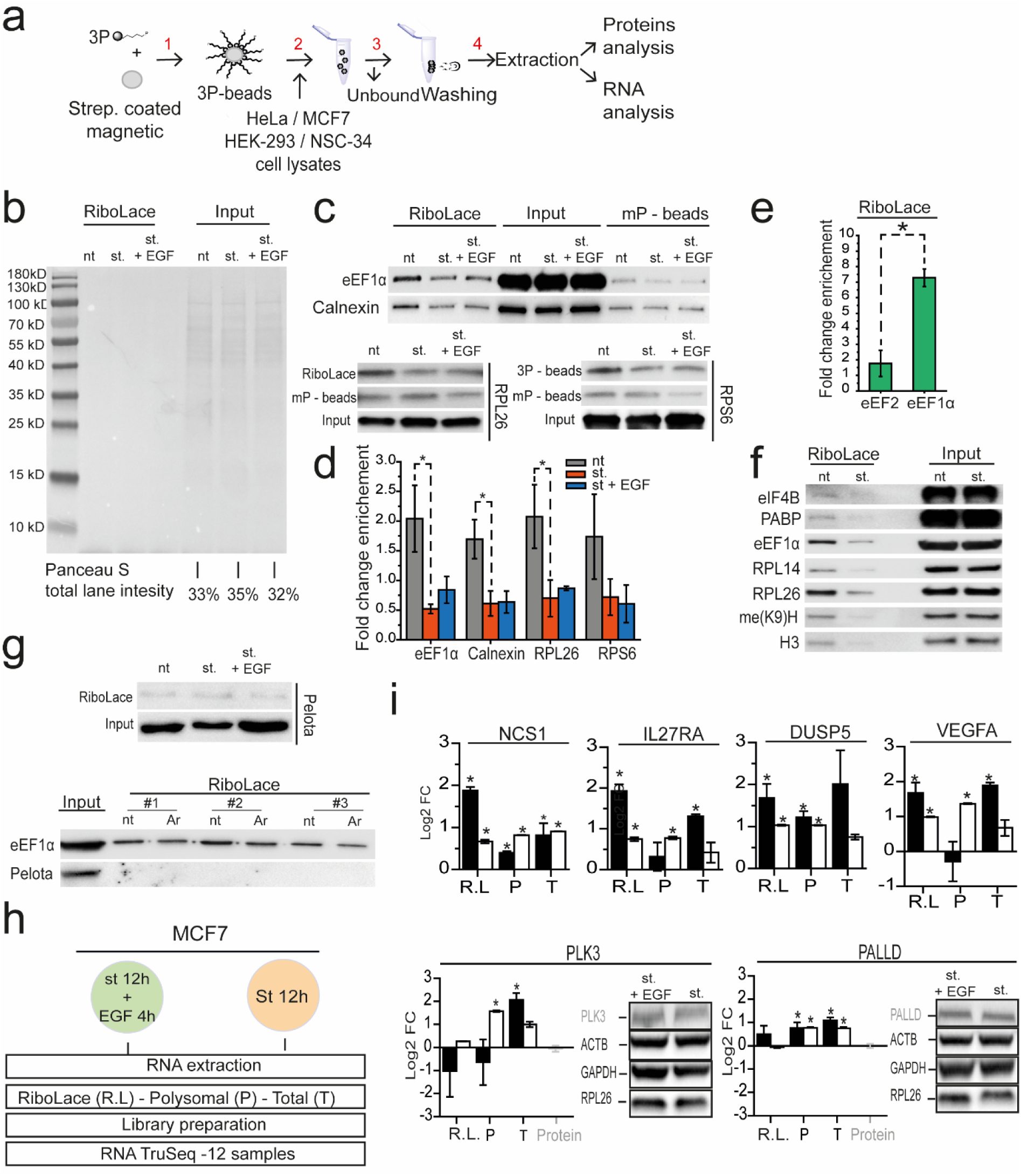
RiboLace is able to capture ribosomes and associated mRNAs under active translation in cell cultures and tissues. **(a)** RiboLace protocol: magnetic beads coated with streptavidin are functionalized with the 3P molecule (step 1). RiboLace beads are then added to the cell lysate (step 2) (usually 5 - 20 μL, corresponding to ∼ 1.2 - 5 x 10^5^ cells) and washed (step 3). Finally, both proteins and RNA are recovered for further analysis (step 4). **(b)** Ponceau staining of a nitrocellulose membrane containing (from left to right) the total protein extract after applying the RiboLace protocol on HEK-293 and corresponding inputs in three different conditions (nt, not treated; st, starvation (0.5% FBS); st + EGF, starvation + EGF stimulation). The lane intensity is reported as a percentage of the sum of the three lysates. **(c)** Immunoblotting of eEF1α, calnexin, RPL26 and RPS6 isolated after applying the RiboLace protocol on HEK-293 (RiboLace, mP-beads, and in input). **(d)** Quantification of proteins isolated with RiboLace in different stress conditions applied to the cells. Immunoblots were scanned with a Chemidoc (Biorad) and quantified with ImageJ v1.45s, n = 3, * indicates pval < 0.05. **(e)** Comparison between the relative enrichment (no starvation vs starvation) of eEF2 and eEF1α on RiboLace. (*) t-test p-val = 0.02, n = 4 **(f)** Immunoblotting of eIf4B, PABP, eEF1α, RPL14, RPl26, me(K9)H3 and H3, detected on RiboLace in normal growing conditions (nt) or under serum starvation (st) with relative inputs in MCF7 cytoplasmic lysates. **(g) Top**, Immunoblotting of Pelota and eEF1α detected on RiboLace in HEK-293 not treated, under starvation or after EGF stimulation, with relative inputs. **Bottom**, Immunoblotting of Pelota and eEF1α detected on RiboLace in MCF7 treated (Ar) or not treated (nt) with Arsenite. (#) indicates the number of each biological replicate **(h)** Experimental design for identifying the global RNA repertoire of RNAs associated to RiboLace by RNA-seq. MCF7 cells treated with EGF or serum-starved in comparison to classical polysomal profiling (POL-Seq) and total RNA transcriptomics analysis. After proteinase K digestion, RNA is extracted from RiboLace and mP beads, and from both polyribosomal and total RNA from the same profile. RiboLace, RL; Polyribosomal, P; Total, T‥ Libraries are prepared using the Illumina TruSeq library preparation kit and the sequencing performed with Illumina HiSeq 2000. (**i) Top**, histograms representing RNA-seq (white bars) and RT-qPCR (black bars) fold changes (FC) of four genes (NCS1, IL27RA, DUSP5, VEGFA) t-test p-val < 0.05 (*). **Bottom**, Comparison between protein fold change and RNA fold changes obtained with different methods (RL, P, T). The semi-quantitative analysis of the protein band intensity (n = 3) for PALLD and PLK3 is reported in the histogram. Black bars, RT-qPCR fold change; white bars, RNA-seq fold change; light gray bars, protein fold change. Housekeeping: GAPDH, beta actin; ribosomal protein L26. t-test (*) = p-val < 0.05.

First, we monitored the enrichment on RiboLace of functional and structural markers of ribosomes in lysates of immortalized human cells (HEK-293T). Importantly, we used 2 x 10^5^ cells, representing ∼ 1/30th of what classical polysomal profiling approaches require. We monitored the ability of RiboLace to purify proteins known to be associated with the translation machinery (eEF1α, calnexin, RPL26 and RPS6). The elongation factor eEF1α is responsible for the delivery of aminoacyl-tRNAs to the translation machinery and is associated to ribosomes in active translation^54,55^. Calnexin is a chaperone protein in the endoplasmic reticulum that associate with ribosomes, helping protein folding during translation^56^. Finally, RPL26 and RPS6 are ribosomal proteins belonging to the large and small subunits of the ribosome, respectively. After immunoblot on the RiboLace eluted proteins, in lysates from untreated cells (nt, Fig 3b and Fig. 3b), we observed an enrichment of all four proteins (Fig. 3c and Fig.3d) with respect to serum starvation (st). When cells were stimulated with EGF after starvation (st. + EGF 1 μg/mL, 4 hours, after starvation), we observed a slight increase in the signal of the translational markers (Fig. 3c). Since the background signal in controls (mP-beads) was the same in all conditions, our results suggest that RiboLace can pull-down active ribosomes and therefore monitoring the translational state of cells.

To further confirm this result, we compared the relative abundances on RiboLace of eEF1α and eEF2 between no-stress and stress conditions. It is known that eEF2-mediated translocation, as well as the switch of ribosome conformation from the non-rotated to the rotated state^57^, is inhibited by cycloheximide^58^. We observed that the two proteins differentially co-sedimented along the polysomal profile, with eEF1α mainly detected in the heavy fractions (**Supplementary Fig. 10b)** indicating a preferential association of eEF1α to ribosomes in cycloheximide treated cells. Then, in agreement with the hypothesis that RiboLace captures active ribosomes, we found enrichment of eEF1α, but not of eEF2 (Fig. 3e and **Supplementary Fig. 10c and Fig 11)**, suggesting the capture of the pre-translocation complex in the non-rotated conformation.

Next, we tested RiboLace in a different cell line, the widely used human tumor cell line (MCF7), in control (nt) or starved (st) and control conditions, again establishing levels of translational markers associated to RiboLace. In addition to ribosomal proteins, we detected additional proteins known to be associated to polysomes (PABP, eIF4B), or a marker of active translation (me(K9)H3)^59^ (Fig 3f and **Supplementary Fig. S10d)**. In agreement with results obtained for HEK-293T cells, the decrease in translational markers associated to the beads in starved cells suggests that RiboLace captures less ribosomes when global translation is downregulated in different cell lines. To further validate this finding, we applied other stresses known to elicit repression of global protein synthesis (i.e. proteotoxic stress, heat shock and sodium arsenite). In all cases, translation markers were reduced (**Supplementary Fig. 10e**). We then tested RiboLace on a mouse motor-neuron like cell line, NSC-34, in normal growth conditions. Also in this case we observed an ∼8-fold enrichment of RPL26 and ∼4-fold enrichment of RPS6 with respect to control beads (**Supplementary Fig. 10f, right)**, demonstrating that RiboLace can efficiently capture ribosomes from both human and mouse cell lines. Lastly, we wanted to establish whether RiboLace could capture components of the eukaryotic surveillance mechanisms that target stalled elongation complexes. Among them, Pelota (mammalian orthologue of the yeast Dom34)^60,61^ is a protein known to promote the dissociation of stalled ribosomes. Strikingly, in control, starved or arsenite treated lysates of HEK-293T and MCF7 cells, Pelota was not enriched on RiboLace beads (Fig. 3g).

These findings prompted us to investigate whether RiboLace can provide an improved estimation of translation efficiency, with respect to the use of total RNA or polysomal RNA in profiling experiments (Fig. 3h and **Supplementary Fig. S12**). We coupled RiboLace to Next Generation Sequencing (NGS) and identified sets of differentially expressed mRNAs associated to RiboLace before and after EGF stimulation (**Supplementary Fig. 13 and Supplementary Fig. 14**). We compared these data with transcripts identified by classical transcriptome (RNA-Seq) or translatome (POL-Seq, i.e. polysomal profiling^62^) analyses. Eight found differentially expressed genes were selected for validation by RT-qPCR (PALLD, PLK3, IL27RA, NCS1, VEGFA, DUSP5, PDCD4, PAPSS2), showing a good agreement with NGS (Pearson’s r = 0.89) **(Supplementary Fig.15)**. Then, we compared the protein levels of four genes (PALLD, PLK3, hb-EGF and CYP27A1) to the relative RNA abundances. PALLD and PLK3 proteins, evaluated by immunoblotting and normalised for three different housekeeping proteins (ACTB, GAPDH and RPL26), did not change upon EGF treatment (Fig 3i). Importantly, only RiboLace (RL) did not show significant variation on both RT-qPCR and RNA-Seq measurements. The same concordance between protein level and RiboLace was obtained with hb-EGF and CYP27A1, whose protein levels increased coherently with RiboLace **(Supplementary Fig. S16)**. Overall, these results establish the important proof-of-concept that RiboLace can capture ribosomes under active translation and determine protein variations more precisely than total or polysomal profiling, at least in our case study.

### *In vivo* active-ribosome profiling using RiboLace

Given the relative simplicity of the RiboLace protocol, and its ability to be used with low amounts of input material, we next wanted to combine our purification strategy with Ribo-Seq. This would establish whether RiboLace could be used to capture active ribosome dynamics along transcripts, improving *in vivo* ribosome profiling experiments. To facilitate this, we modified our original protocol by including an endonuclease digestion step and applied it to lysates from mouse tissues (Fig. 4a).

**Figure 4.**
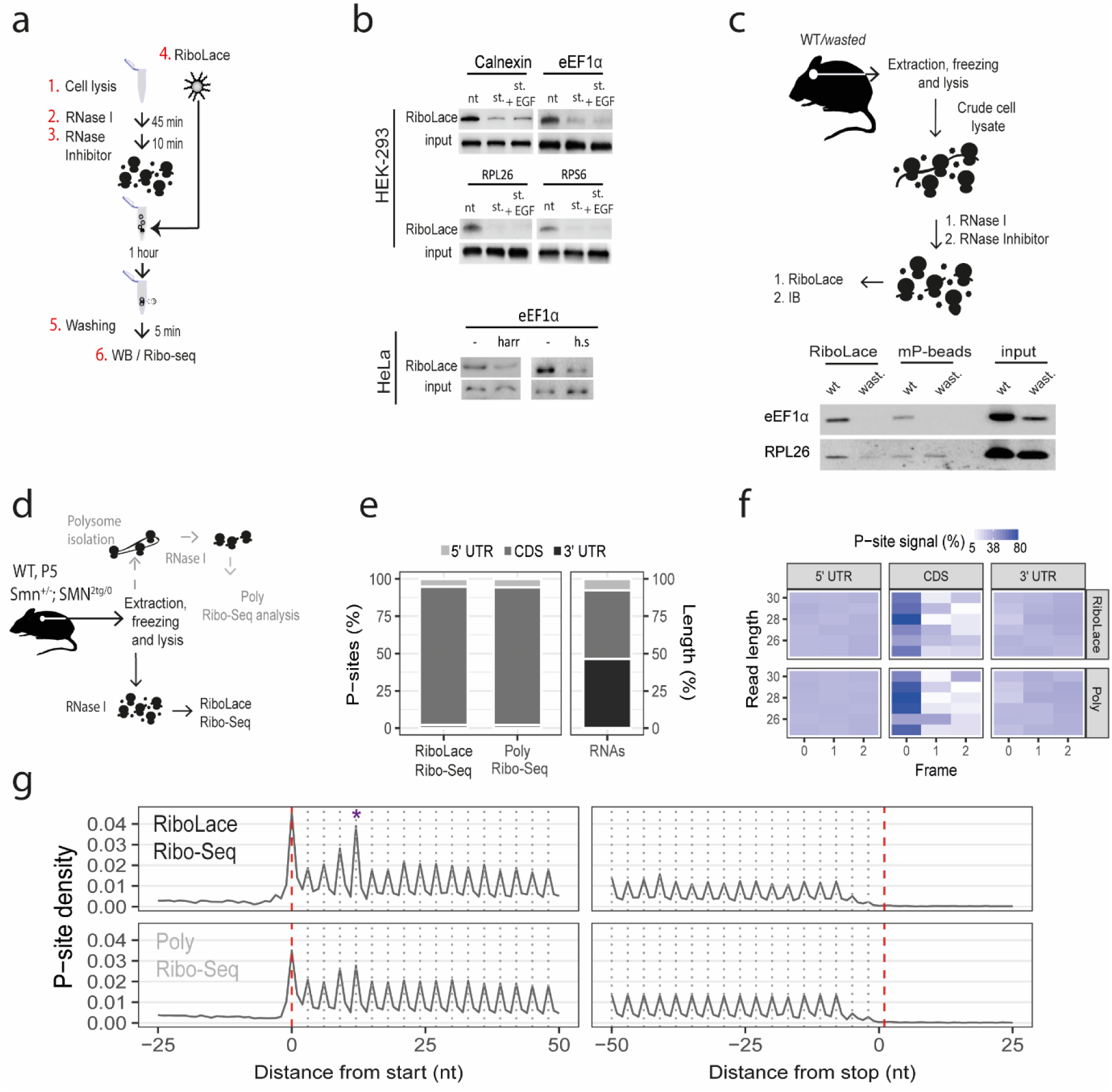
RiboLace Ribo-Seq. **(a)** Schematic RiboLace protocol for the separation of single active ribosomes. Cell lysates (1) is treated with RNase I for 45 min (2) and the quenched with an RNase inhibitor (3). Then, RiboLace beads are incubated with the digested cell lysate (1h, 4°C) (4), washed (5) and then proteins or RNA extracted (6). See methods for details. **(b)** Immunoblotting of eEF1α, calnexin, RPL26 and RPS6 after applying the RiboLace protocol on HEK-293 and HeLa. Nt, not treated; st, starvation (0.5% FBS), st + EGF, starvation + EGF stimulation; harr, harringtonine; h.s., heat shock **(c) Top**, Schematic RiboLace protocol for the extraction of single active ribosomes from mouse brain. **Bottom**, immunoblotting on eEF1α, and RPL26 for RiboLace, mP-beads and relative inputs. **(d)** Scheme of the protocol for comparative active Ribo-Seq and poly Ribo-seq. **(e) Left**, percentage of P-sites mapping to the 5’ UTR, CDS and 3’ UTR of mRNAs from RiboLace and Poly Ribo-Seq data. **Right**, percentage of region lengths in mRNAs sequences. Both techniques show a clear enrichment in signal mapping to the CDS, consistent with ribosome protected fragments. **(f)** Percentage of P-sites corresponding to the three possible reading frames along the 5’ UTR, CDS and 3’ UTR, stratified for read length, comparing RiboLace (top panel) and Poly Ribo-Seq (bottom panel). For each length and each region, the sum of the signal is normalized to 100%. The enrichment in frame 0 is CDS specific in both cases. **(g)** Meta-gene profiles showing the density P-sites around the translation initiation site (TIS) and translation termination site (TTS) for RiboLace (top panel) and Poly Ribo-Seq (bottom panel). The peak corresponding to the fifth codon is highlighted with an asterisk.

To demonstrate that RiboLace can capture isolated ribosomes after endonuclease digestion, we first measured the enrichment of eEF1α, calnexin, RPL26 and RPS6 on RiboLace applied to control (nt), harringtonine treated (harr), serum starved (st) or heat-shocked (h.s.) HEK-293T and HeLa cells (Fig. 4b). Our results confirmed that RiboLace can selectively capture isolated ribosomes under conditions of active translation. Then, we confirmed that RiboLace was able to enrich ribosome protected fragments, by urea-gel electrophoresis and by the use of the Bioanalyzer (**Supplementary Fig. S17**). After that, we probed RiboLace on 15 μL (∼ 1/50thof the total lysate) samples of whole brain lysates from wild-type (WT) and mice affected by the “*wasted”* mutation, consisting in a deletion of the elongation factor eEF1α2 **(Fig.4c)**, resulting in defective translation^63^. Our results, in agreement with those previously obtained from cells, showed that RiboLace could selectively enrich RPL26 only in the WT mouse tissue, extending its functionality from cell lines to tissues.

Having established that RiboLace can capture isolated ribosomes from very low amounts of starting material, we next isolated ribosome protected fragments 25-35 nt long (RPFs) from ribosomes pulled down by RiboLace and, in parallel, from polysomes isolated from whole mouse brain, following nuclease digestion (polysomal Ribo-Seq, Fig. 4d and **Supplementary Fig S18**). After sequencing, we analyzed both RiboLace and polysomal Ribo-Seq RPFs using the dedicated pipeline RiboWaltz^64^ to obtain sub-codon information and identification of the trinucleotide periodicity. The distribution of the reads showed a main population of length at ∼28 nt (**Supplementary Fig. S19**), in agreement with what has previously been observed for ribosomes trapped on the mRNA by cycloheximide^65,66^. As expected for ribosome footprints, we observed an enrichment of signal along the coding sequence in both RiboLace and polysomal Ribo-Seq data (Fig. 4e). Occupancy meta-profiles showed the typical trinucleotide periodicity of the ribosome P-site along coding sequences, which is suggestive of signal derived from translating ribosomes (**Fig.4f and g**). The comparison between meta-profiles obtained with RiboLace and polysomal Ribo-Seq highlights an accumulation of ribosomes at the start codon and at the 5^th^ codon (Fig. 4h). Interestingly, the latter feature is associated with ribosome pauses necessary for a productive elongation phase of translation^67^.

Taken together, these results confirm that RiboLace is capable of providing positional data with nucleotide resolution and of enriching samples with active ribosomes, thereby facilitating reliable descriptions of *bona fide* translational events *in vitro* and *in vivo*.

## DISCUSSION

During their lifetime in the cellular cytoplasm, mRNAs are regularly stored, degraded, and transported, with only a fraction being actively translated to produce proteins^68,69^. All these stages of the mRNA lifecycle are governed by *cis*- and *trans*-factors that tightly regulate the uploading of mRNAs on polysomes, and subsequent production of proteins. In order to generate a better understanding of these sophisticated and dynamic processes, different methodological approaches have been developed to determine, at a genome-wide scale changes in RNA steady state levels (e.g. RNA-seq), the change in engagement with the translational machinery (e.g. Ribo-Seq, polysomal profiling)^20,62^, and the change in protein production (e.g., SILAC, PUNCH-P)^40,70,71^. Although Ribo-Seq remains a complex technology that requires relatively large tissue samples, it has been shown to be extremely powerful for identifying ORFs and translation initiation sites (i) from cell lysates or ribosome pellets; (ii) from purified polysomal fractions (polysomal Ribo-Seq)^19^ or, more recently, (iii) from tagged ribosomes^72,73^. Unfortunately, however, the use of cell lysates and ribosome pellets often introduces unwanted background signals, presumably mostly due to the presence of stalled ribosomes and fragments protected by RNPs, or by the 80S monosome.

Here, we sought to significantly enhance Ribo-Seq approaches by developing a new molecule (3P) that facilitates the selective capture of ribosomes under active translation. We focused our attention on puromycin, the well-known structural analogue of the 3′ end of aminoacyl-tRNA, and we suppressed its irreversible activity by tethering the α-amino group to a biotinylated linker. Despite this modification on the primary amino group of puromycin, we observed that 3P could still interfere with eukaryotic translation *in vitro.* We used 3P-functionalized magnetic beads (a method we called RiboLace) to capture and enrich transcripts undergoing translation in eukaryotic *in vitro* and *in vivo* systems. We observed that the elongation factor eEF1α, a key protein involved in delivering tRNAs to the ribosome, was the most enriched protein on RiboLace in all our experiments. This may be explained with the binding of 3P to the A-site of the ribosome in the not-rotated state, when the acceptor site accommodates the aminoacyl-tRNA engaged by eEF1α. We then demonstrated that RiboLace is capable of providing positional data with nucleotide resolution of translational events when used for ribosome profiling. Remarkably, we observed > 95% of RPFs on the coding region with the characteristic trinucleotide periodicity, suggestive of active ribosomes flowing along the transcripts. In our biological model, RiboLace Ribo-Seq showed almost no signal on the 5′- and 3′-untranslated regions of the mRNAs, and a peculiar pause of ribosomes at the 5^th^ codon. Overall, our data suggest that RiboLace possess significant advantages, in terms of both sample yield and accuracy in RPFs detection, over classical Ribo-Seq and polysomal Ribo-Seq approaches.

RiboLace protocols can be further adjusted to (i) isolate ribosomes from other organisms, (ii) isolate ribosomes from specific eukaryotic cellular compartments such as the Endoplasmic Reticulum or organelles like mitochondria, or (iii) to characterized active specialized-ribosomes^2,72,74^ induced by different cell conditions. We specifically designed RiboLace for ribosome profiling experiments, to facilitate improved understanding of ribosome dynamics along transcripts and to allow better estimates of translation levels based on ribosome footprints.

In summary, we report a major advancement in the technology available to undertake ribosome profiling, providing the unique benefit of capturing ribosome in active translation. Given its ability to enrich actively translating ribosome protected fragments, RiboLace can be used in difficult samples with low input material, and empowers accurate conclusions to be drawn concerning the actual translational state of a biological system, paving the way for more detailed and accurate studies of translation in the future.

## METHODS

Methods and Supplementary Figures are available in the file Supplementary Information.

## ACKNOWLEDGMENTS

We thank Tocris Bioscience (a Biotechne brand) for the support in scaling up the 3P synthesis. We thank Divya Kandala and Luca Minati for the helpful discussions. We also thank Veronica Desanctis and Roberto Bertorelli of the CIBIO NGS for technical support and Daniele Arosio for the kindly gift of the IPR-IBA2 plasmid.

## AUTHOR CONTRIBUTION

MC and GV conceived the experiments. MC performed all the biological experiments and contributed to the 3P synthesis. TT, DG and FL analyzed the RNA-seq and Ribo-Seq data. PB and EP performed the Poly Ribo-Seq and helped with the RiboLace Ribo-Seq. LP performed control experiments with mP-beads. RG performed a preliminary synthesis of 3P. GG designed the 3P synthesis, performed MS analysis, COSY and NMR analysis of the 3P. EJNG and THG generated and provided the mouse tissues. MC and GV drafted the manuscript. TT, FL and DG wrote the methods related to RNA-seq and Ribo-Seq data analysis. GG wrote the methods related to the chemical synthesis and characterization. GV, MC, TT, FL, GG, THG and AQ reviewed the manuscript. All authors read and approved the final manuscript.

## Data availability

Raw and analyzed data for Ribosome profiling have been deposited under GEO: GSE102354 for RiboLace Ribo-Seq, and GEO: GSE102318 for Poly Ribo-Seq

